# Development and tissue specific expression of RAPGEF1 (C3G) transcripts having exons encoding disordered segments with predicted regulatory function

**DOI:** 10.1101/2024.04.07.588436

**Authors:** Archana Verma, Abhishek Goel, Niladri Koner, Gowthaman Gunasekaran, Vegesna Radha

**Author notes:** Correspondence: Vegesna Radha, CSIR-Centre for Cellular & Molecular Biology. equal contribution (Archana Verma, Abhishek Goel). Archana Verma, Dept of Pediatric Hematology and Oncology University childrens hospital, Muenster, Germany-48149. Gowthaman Gunasekaran, Department of Molecular Biology Laboratory of Chromatin Biology Ariel University Ariel Israel - 40700.

## Abstract

The ubiquitously expressed RAPGEF1(C3G), regulates differentiation, and is essential for development of mouse embryos. While multiple transcripts have been predicted, evidence of their expression and function is scarce. We demonstrate tissue and development specific expression of novel transcripts with exons 12-14 in various combinations, in the mouse. These exons encode an intrinsically disordered serine-rich polypeptide, that undergoes phosphorylation. Isoform switching occurred during differentiation of myoblasts and mouse embryonic stem cells. In silico structure and docking studies indicated that the additional exons alter intra-molecular interactions keeping it in a closed confirmation, and interaction with its target, RAP1A. Our results demonstrate the expression of novel RAPGEF1 isoforms, and suggest cassette exon inclusion as an additional means of regulating RAPGEF1 activity during differentiation.

## Introduction

Cell fate determination and tissue specification during development are regulated by external signals and their transducers, which control gene expression and cytoskeletal reorganization. RAPGEF1(C3G), a guanine nucleotide exchange factor (GEF) is essential for early embryonic development of mice [1], and has recently been identified as a determinant of the balance between proliferation & differentiation of mouse embryonic stem cells (mESCs) [2]. Absence of RAPGEF1 in these cells resulted in enhanced self-renewal and an inability to differentiate. It is a ubiquitously expressed protein involved in signalling from growth factors, cytokines, and adhesion receptors, by activating small GTPases of the Ras family [3]. In addition, RAPGEF1 can engage in signaling through multi-molecular complex formation [4, 5]. It is a protein of 140kDa, with a catalytic domain at its C-terminal, a central region with multiple poly-proline tracts involved in protein interaction, and an N-terminal containing an E-cadherin binding domain. Catalytic activity of RAPGEF1 is regulated through intra-molecular interactions, phosphorylation, and membrane localization [3]. Its activity is kept in check by an auto-inhibitory region (AIR), present within the Crk binding region (CBR), that folds over and inhibits GEF activity [6]. Residues in the N-terminal interact with Ras exchanger motif within the catalytic domain to enhance its activity. RAPGEF1 interacts with and is phosphorylated by Src family kinases, Abl, and GSK3β, which regulate its function, and sub-cellular localization[7–9]. It also forms complexes with Crk, Cas, β-catenin, cenexin, and TC-PTP[4, 10–13]. RAPGEF1 is engaged in cell fate determination through modulation of gene expression, survival signaling, centrosome and primary cilium dynamics and, cytoskeletal remodelling[3, 13–15]. Its deregulation is associated with multiple disorders[16–19].

RAPGEF1 is expressed from a single gene located on chromosome 9 in humans, and chromosome 2 in mouse, and shows high degree of homology across vertebrate species. In the mouse, 27 transcripts have been predicted, of which four are protein coding and some evidence is available for their expression in a development and tissue specific manner [20–22]. Despite an understanding of its role in multiple signaling pathways, evidence for expression of various isoforms, and their function in embryonic and adult mouse tissues is scarce. Understanding the properties of the various isoforms, and the specific role they play during development and in function and identity of adult tissues has been challenging, but of considerable interest. In an attempt to study the expression patterns of RAPGEF1 isoforms, we examined embryonic and adult mouse tissues, and various cell lines by splicing specific-PCR assays. We demonstrate the specific expression of isoforms of varying length in adult tissues like the brain, heart, testis, and skeletal muscle, all of them showing unique patterns due to insertion of one or more cassette exons, at a hot spot adding additional amino acids between the protein interaction domain and catalytic domain. We also show isoform switching during differentiation of myoblasts into myotubes. The 3D structures of RAPGEF1 isoforms were predicted using AlphaFold [23, 24], and protein-protein docking with RAP1A was examined using LZerD [25, 26]. Our analysis identified the contribution of cassette exons to autoregulation of its catalytic activity, and differences in ligand interaction. Data suggest that the products of alternately spliced isoforms of RAPGEF1 are regulated by cis-acting domains present in some of them.

## Materials & Methods

### Materials

cDNA preparation kit, and DNA size marker ladders were from Takara. TRIzol reagent from Thermo Fisher Scientific was used for RNA isolation. EmeraldAmp 2X PCR Master Mix was used for performing all PCRs. DMEM, FCS, and Horse serum, were from GIBCO. Reagents used for SDS-PAGE & Western blotting were procured from Sigma. GFP antibody was from Santa Cruz (sc8334). Primer synthesis was by Reprocell.

### Cell lines, culture and treatments

HL-1 mouse cardiomyocyte cell line was procured from Millipore (Cat. # SCC065), and grown as indicated by the supplier. C2C12, mouse myoblast cell line was a gift from Dr. Helen Blau, and was grown and differentiated as described [14]. Photoreceptor cell line 661W was provided by Dr. Muyyad R. Al Ubaidi [27]. E14Tg2A cells were grown in hanging drop culture for 48 h and embryoid bodies were transferred to media without LIF to facilitate differentiation [2]. All other cell lines were obtained from ATCC, and grown in DMEM with 10%FCS, in the absence of antibiotics. Okadaic acid (OA) and pervanadate (PV) treatments were carried out at 10nM for 8hrs; and 50micromol for 10mins, respectively.

### Animal and Tissue Processing

The work was carried out after approval by the Institutional animal ethics committee (IAEC) of CCMB, no. IAEC 08/1017, and protocols strictly follow the guidelines of Committee for the purpose of control and supervision of experiments on animals (CPCSEA), Government of India for animal welfare. Wild type male & female C56 black mice of 8 weeks, and 12months age, were used in this study. Embryonic tissues were collected at 18dpc. Uterus and whole embryo were collected at 8dpc. Different tissues were collected in liquid N2, with the help of an authorised technical officer, and stored frozen in −70 freezer.

### RNA isolation, cDNA Preparation, primer designing, Semi-quantitative PCR and sequencing

Cells at 75% confluency in a T25 flask and 50-100 mg tissue samples were used for RNA isolation using the TRIzol method Rio DC, 2010). RNA samples were subjected to cDNA synthesis from 1μg of RNA using Takara Bio cDNA synthesis kit and stored at −20°C. The RAPGEF1 mouse isoforms 1 & 2 (NM_001039087.1 & NM_001039086.1) were used as a template for designing the primers using Primer3Plus tool and validated by blasting in NCBI.

PCR was carried out using the prepared cDNA for amplification of RAPGEF1 isoforms and GAPDH. List of primers used and their sequence is shown. For GAPDH amplification; initial denaturation was at 98°C for 2 mins followed by 25 cycles of denaturation for 15 sec at 98°C, annealing at 58°C for 30 sec, and extension at 72°C for 30 secs. and a final extension for 5 mins at 72°C. Similar conditions were used for the N-term primers, except that amplification was carried out for 35 cycles. The following PCR conditions were used for amplification of isoforms, with and without exons 12, 13 & 14. (RAPGEF1 isoF, forward primer, and one or other reverse primers): initial denaturation was at 98°C for 30 sec, followed by 35 cycles of denaturation at 98°C for 15 sec; annealing at 62°C for 30 sec. and final extension for 5mins at 72°C. Sequencing was done at the sequencing facility at CSIR-CCMB.

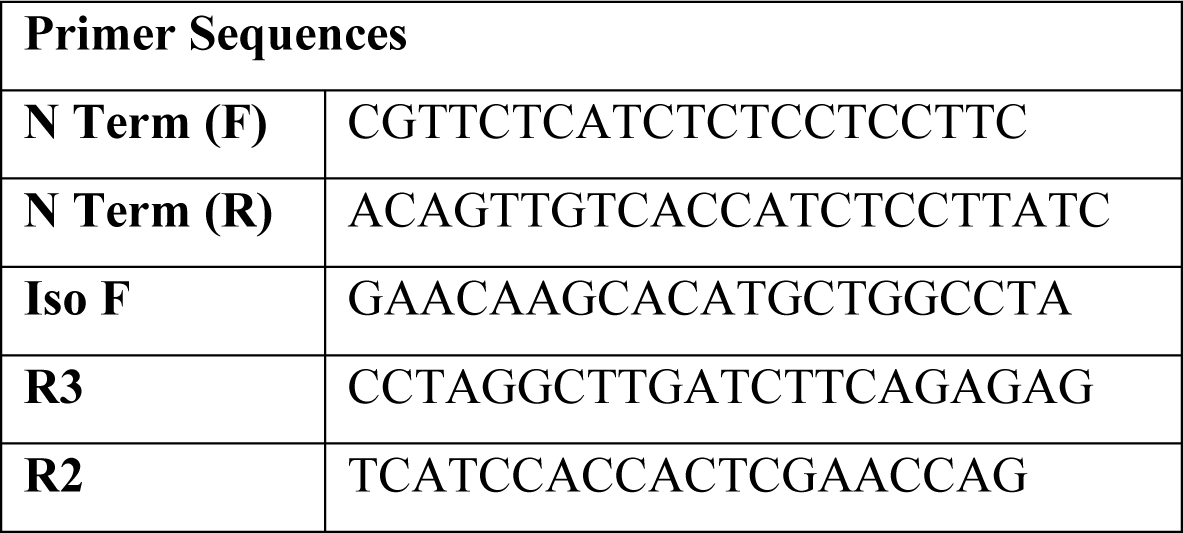
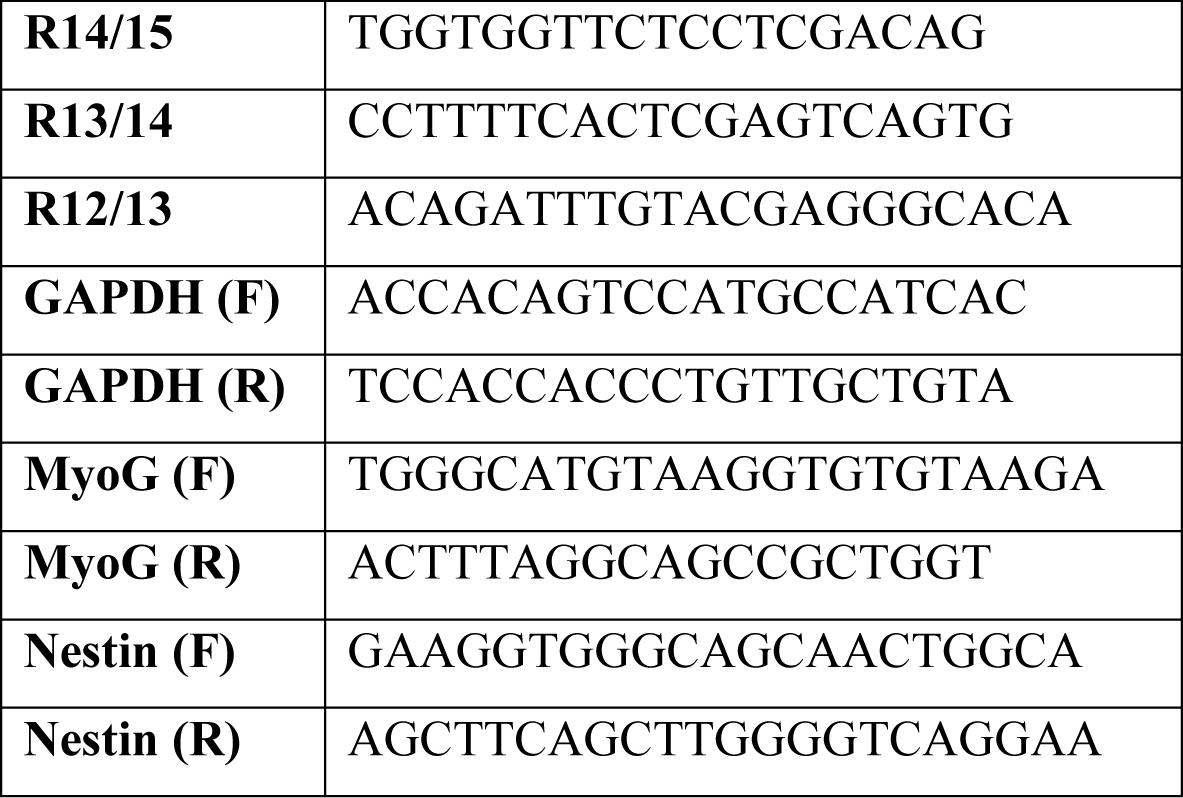

### Generation of vector expressing exons 12 & 13 as a GFP fusion protein

The 414 bp insert in mouse RAPGEF1 isoform 1 was amplified from cDNA prepared from mouse brain RNA, and cloned into pEGFP-N3 vector, using the primers as indicated (Target sequence 2130-2614 of mouse isoform 1) using Xho1 and BamH1 restriction sites.

RAPGEF1 INSERT GFP-N3 1224 FP:

5’ – TAAGCACTCGAGATGCCACCCGCTCTGCCCCCCAAGCAACGGCAGC, and RAPGEF1 INSERT GFPN3 1224 RP

5’ – TGATTGGGATCCGGGATGTCCGTCCTTCCCAGATGGTGGTTCC Clones with the inserts were confirmed by sequencing.

### SDS-PAGE and Western blotting

Lysates from cells grown in culture and mouse tissues were prepared and subjected to western blotting as described [22].

### Structural studies

NetSurfP - 3.0 – tool was used for Protein secondary structure and relative solvent accessibility determination. 3D models of mouse RAPGEF1 and mouse RAP1A were accessed from the AlphaFold protein structure database (https://alphafold.ebi.ac.uk<u>)1</u>. PyMOL visualisation tool was used to investigate the intramolecular interactions of three significant residues found in the AIR region: M523, Y526, and M527. For Proximity Operations, the “around” command was used with a cutoff distance of “within 5 Ang” which displayed all catalytic domain residues (from three of the helices, CD α1, CD α2, and CD α3) within 5 Ang surrounding AIR residues. To analyze the interaction between the different isoforms of mouse RAPGEF1, and mouse RAP1A, RAPGEF1 isoform1 (Q3UHC1_MOUSE, NP_001034176.1, aa 540 to 1224), RAPGEF1 isoform2 (Q3UGX8_MOUSE, NP_001034175.1, aa 502 to 1218), RAPGEF1 isoform3 (Q91ZZ2_MOUSE, NP_473391.1 aa 540 to 1086), and full length RAP1A (P62835) were used for docking analysis in order to save computational resources. Unbiased rigid body docking (exhaustive search of all possible binding sites and binding poses) was performed using LZerD Web Server3, pairwise protein docking (https://lzerd.kiharalab.org/upload/upload/). For the docking, we set no constraints and default cluster cutoff (root-mean square deviation, RMSD, of 4 Å). We selected as binding sites all the regions where the number of contacts with RAP1a for each residue was greater than 600. After calculating the distance between carbon α of the interacting residues, the best docked pose was detected according to the MM/GBSA sever, based on the free binding energy (−18.11 kcal/mol for RAPGEF1 isoform1, −37.57 kcal/mol for RAPGEF1 isoform2 and −23.3 kcal/mol for RAPGEF1 isoform3). The visual representation of models and docking results were generated by the software package BIOVIA Discovery Studio Visualizer4.

## Results

### Tissue specific expression of novel alternately spliced isoforms of mouse

Located on the positive strand of chromosome 2, mouse RAPGEF1 spans 120.6 kb, and the current annotation release from NCBI, indicates 27 exons, and 27 transcript variants, with the expression of four protein coding isoforms being confirmed. These isoforms along with schematic showing domain organization of RAPGEF1 protein are indicated in Fig. 1A, B & Table 1. Apart from the start site, the various isoforms differ particularly in the inclusion or skipping of exon 3, and exons 12,13, and 14, which are cassette exons. Isoform 3 lacks exons 12, 13 & 14, but has exon 3. Isoform 1 has exons 3, 12 & 13, while isoform 2 lacks exon 3, but has the whole cassette of exons 12, 13 & 14. Isoform 4 is similar to isoform 2, but lacks exon 3. Positions of the variably expressed exons and the number of amino acids they encode are indicated in Fig. 1B, and the amino acid alignments of the three full length isoforms is shown in Fig. S1.

**Table: 1.** List of experimentally identified and predicted full length RAPGEF1 transcripts indicating presence or absence of exon 3, and the cassette exons 12, 13, and 14. The length of the transcripts, and the corresponding proteins are indicated.

| NCBI Transcript ID | Ensembl Transcript ID | NCBI Transcript name | NCBI Protein | Length (nt) | Length (aa) | Exon 3 | Cassette exons |
| --- | --- | --- | --- | --- | --- | --- | --- |
| XM_006497583.2 |  | X2 | XP_006497646.1 | 6348 | 1256 | + | 12,13,14 |
| XM_030246484.1 |  | X17 | XP_030102344.1 | 5994 | 1138 | + | 12 |
| XM_036157277.1 |  | X4 | XP_036013170.1 | 6345 | 1255 | + | 12,13,14 |
| XM_036157284.1 |  | X8 | XP_036013177.1 | 6249 | 1223 | + | 12,13 |
| XM_036157298.1 |  | X13 | XP_036013191.1 | 6138 | 1186 | - | 12,13 |
| XM_030246482.1 |  | X11 | XP_030102342.1 | 6231 | 1217 | - | 12,13,14 |
| XM_030246487.1 |  | X22 | XP_030102347.1 | 5724 | 1048 | - | - |
| XM_030246485.2 |  | X19 | XP_030102345.1 | 6097 | 1127 | - | 12,13,14 |
| XM_030246488.2 |  | X23 | XP_030102348.1 | 5743 | 1009 | - | 12 |
| NM_001039087.1 | ENSMUST00000095087<br>(Rapgef1-202) | 1 | NP_001034176.<br>1 | 6251 | 1224 | + | 12,13 |
| NM_054050.2 | ENSMUST00000102872<br>(Rapgef1-203) | 3 | NP_473391.1 | 5837 | 1086 | + | - |
| NM_001039086.1 | ENSMUST00000091146<br>(Rapgef1-201) | 2 | NP_001034175.<br>1 | 6233 | 1218 | - | 12,13,14 |
| NM_001362702.1 | ENSMUST00000147755<br>(Rapgef1-207) | 4 | NP_001349631.<br>1 | 6134 | 1185 | - | 12,13 |
| XM_036157280.1 |  | X6 | XP_036013173.1 | 6222 | 1225 | + | 12,13,14 |
| XM_006497582.4 |  | X1 | XP_006497645.1 | 6286 | 1257 | + | 12,13,14 |
| XM_006497585.5 |  | X5 | XP_006497648.1 | 6190 | 1225 | + | 12,13 |
| XM_006497589.4 |  | X16 | XP_006497652.1 | 5932 | 1139 | + | 12 |
| XM_006497584.4 |  | X3 | XP_006497647.1 | 6283 | 1256 | + | 12,13,14 |
| XM_030246481.2 |  | X7 | XP_030102341.1 | 6187 | 1224 | + | 12,13 |
| XM_006497590.4 |  | X18 | XP_006497653.1 | 5929 | 1138 | + | 12 |
| XM_006497587.5 |  | X12 | XP_006497650.1 | 6130 | 1205 | + | 13,14 |
| XM_036157299.1 |  | X15 | XP_036013192.1 | 6034 | 1173 | + | 13 |
| XM_006497592.4 | ENSMUST00000238899<br>(Rapgef1-208) | X20 | XP_006497655.1 | 5776 | 1087 | + | - |
| XM_006497586.5 |  | X9 | XP_006497649.1 | 6172 | 1219 | - | 12,13,14 |
| XM_036157288.1 |  | X10 | XP_036013181.1 | 6169 | 1218 | - | 12,13,14 |
| XM_030246483.2 |  | X14 | XP_030102343.1 | 6073 | 1186 | - | 12,13 |
| XM_036157304.1 |  | X21 | XP_036013197.1 | 5662 | 1049 | - | - |

**Fig. 1:**
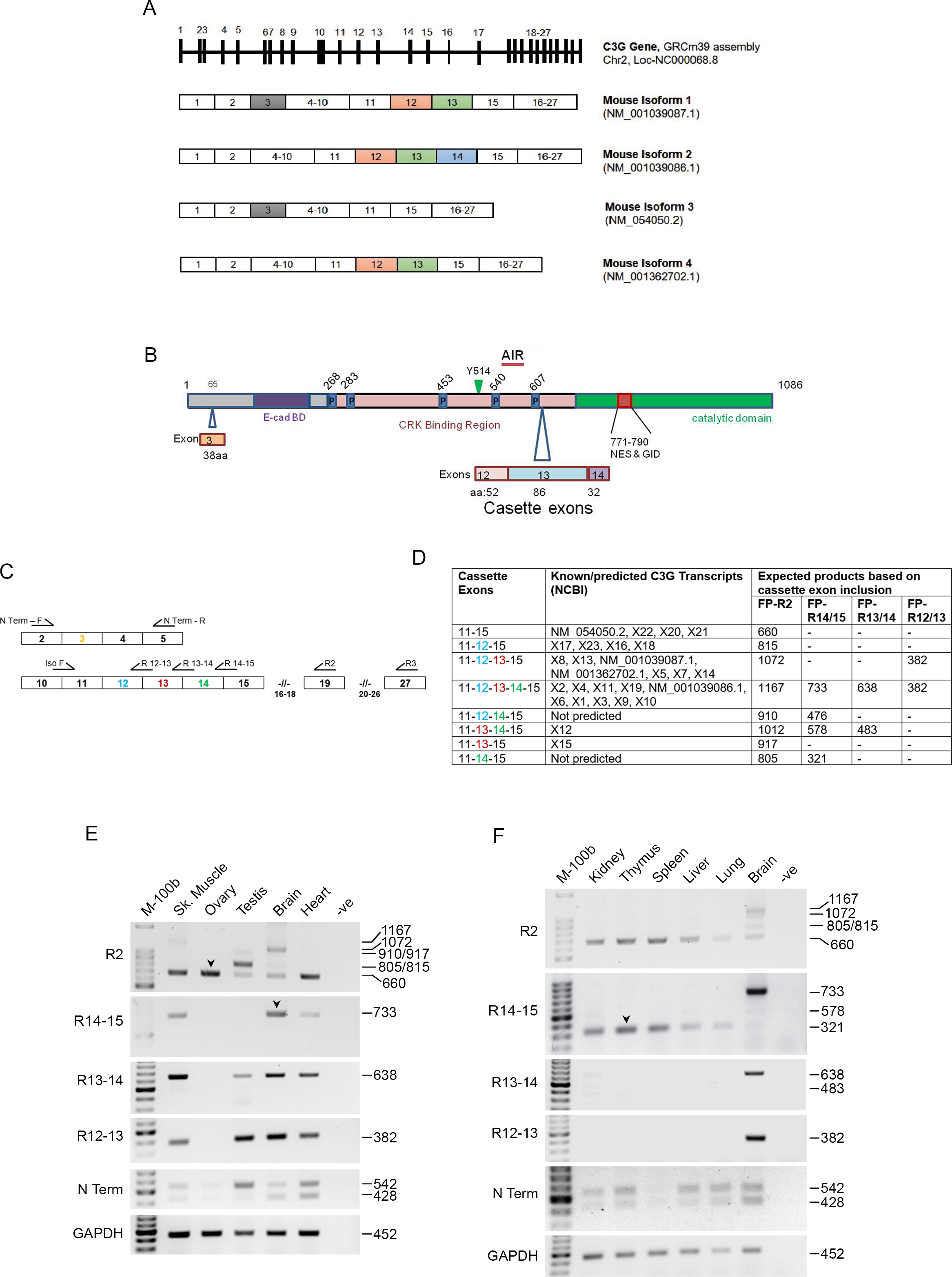
Expression of alternately spliced isoforms of RAPGEF1 in mouse tissues. A) Schematic showing *MmRAPGEF1* gene indicating positions of the 27 exons. Transcripts of the four experimentally confirmed protein coding isoforms with their exons are shown below. They primarily differ in splicing of exon 3, and the cassette exons, 12,13, & 14. B) Schematic showing domain organization of RAPGEF1 protein indicating location of insertion of segments (with number of residues indicated) by cassette exons. The amino acid position numbering given is for isoform 3, which includes exon 3. P, poly-proline tract; AIR, auto-inhibitory region; NES, nuclear export sequence; GID, GSK3β interaction domain. C) Schematic showing positions of primers used to identify transcripts with inclusion or exclusion of exons 3, 12,13, & 14. D) Table indicating expected product sizes with each of the primer pairs. E & F) Amplicons obtained upon PCR of the mouse tissues amplified using the indicated primers. The two products obtained using N-term primers are those with and without exon 3. Product sizes are indicated on the right side. GAPDH was used as internal control. Arrow heads indicate the amplicons that were sequenced for confirmation.

We examined the expression of RAPGEF1 isoforms with inclusion of exon 3, and 1 or more of the other cassette exons. Primers were designed to amplify cDNAs generated from various mouse tissues that would give rise to different product sizes. These are indicated in Fig 1C & D. RNA was isolated from various embryonic and adult mouse tissues, as well as a few cell lines, and cDNAs generated by reverse transcription (RT). The products were used as templates in PCR reactions using N-term primers that generate amplicons with inclusion or exclusion of exon 3 (542 & 428bp). GAPDH was amplified as an internal control. As shown in Fig.1E &F, all tissues from 8wk old mice express isoforms that retain or skip exon 3. While some tissues express equal amounts of these products, others show difference in relative amounts of the two types of products, with the testis predominantly expressing isoforms with exon 3.

Multiple primer pairs were used to detect expression of transcripts with cassette exons, 12,13, and 14 (primer sets, F-R2, F-R14-15; F-R13-14, and F-R12-13). The primers were designed to match exon junctions, to avoid non-specific amplification. R2 would give products of varying lengths depending on which of the three cassette exons are included in the transcripts. PCR was performed under saturating conditions of 35 cycles to detect minor products. As a negative control, RNase free water was used instead of the template. As described earlier, isoform 3, which lacks the cassette exons is the predominant isoform expressed in most tissues and cell lines, and gives amplicons of 660 bases when amplified using primer R2 (Fig.1E & F). This is the transcript that encodes the 140kDa polypeptide. Kidney, Thymus, spleen, liver, ovary and lung, predominantly express isoform 3. The 660bp product obtained from the ovary was sequenced to confirm the contiguity of exon 11 with exon 15 (Fig. S2A). Few tissues, like the brain, heart, testis and skeletal muscle showed the presence of additional amplicons indicative of the inclusion of one or more cassette exons. Each of these tissues had a signature pattern of isoform expression, with 1072bp product being most predominant in the brain (isoform 1), and 805, or 815bp product being predominant in the testis (novel isoform), when examined using the R2 primer. Alternate minor products of 660, 910/917, and 1167 bases are also seen in these tissues. The heart and skeletal muscle show additional longer products, though 660 amplicon is the major form. Eight alternate transcripts (and amplicons) can be predicted to arise with inclusion of the three cassette exons in various combinations, but only three amplicons of 660, 1072 and 1167 were expected from the previously identified transcripts (Fig. 1D, and Table 1), and correspond to isoforms 3, 1 or 4, and 2 respectively. The alternate sizes of the PCR products obtained suggested that the cassette exons are being expressed in various combinations, some of them previously neither identified, nor predicted. The alternate isoforms were predominantly expressed in highly differentiated tissues, where majority of cells do not proliferate, like the brain, heart, skeletal muscle and testis.

An additional reverse primer R3 corresponding to exon, 27, that was used earlier to identify the brain specific isoform [22], was also used to examine amplicon lengths, and compared with products obtained with R2 primer located in exon 19. The sizes of the amplicons obtained with these primers corresponded with the predicted sizes of products with differences only if cassette exons are included suggesting that no other transcript variants are found that differ in any of the exons between 19 and 27 (Fig. S3).

To confirm the presence of each of the cassette exons in these tissues, we used primers that overlap 12-13, 13-14 and 14-15 exons. These primers would not give amplicons from transcripts lacking the cassette exons. The expected amplicon sizes are indicated in Fig.1D. cDNAs from various tissues were subjected to PCR using these primer sets. Brain, testis, heart and skeletal muscle showed products corresponding to isoforms that have the cassette exons in various combinations (Fig 1E & F). The 733bp product from the brain that corresponds to transcripts with all three cassette exons was sequenced for confirmation (Fig.S2B). Interestingly, we found expression of an amplicon of 321bps, which corresponds to a transcript lacking exons 12 & 13, but having exon 14. We sequenced this product amplified from thymus RNA, and confirmed that it as a RAPGEF1 amplicon (Fig.S2C). This is a previously undetected and unpredicted isoform. We also observed predominant expression of an isoform that includes only exon 12, but not 13 and 14, in the testis, as evidenced from the 815bp product with R2 primer, and absence of amplification with 14-15 primer sets. The skeletal muscle, brain and heart express isoforms 1 and 2/4 at different levels. Relative difference in the products was indicative of their differential expression in various tissue types.

Since brain, heart and skeletal muscle showed multiple alternate isoforms, we tested their expression in cell lines, originating from these tissues. Three mouse neuronal cell lines, N2A, 661W, and NSC34; a myoblast line, C2C12, and a cardiomyocyte line, HL1 were examined. All of them showed expression of transcript 3 (Fig 2A). Except HL1, the others showed expression of the novel transcript with only exon 14. Unlike adult heart tissue, HL-1 cells that have properties of differentiated cardiomyocytes, but continue to proliferate, did not show transcripts with any of the cassette exons. The neuronal cell lines lacked significant expression of the cassette exons, with very weak amplicon of 472bp seen in N2A cells, a feature very different from brain tissue. The myoblast cells showed an amplicon of 483bp with R13-14, a product obtained from expression of transcripts with exons 13 & 14, but lacking exon 12. This has been predicted as variant XM_006497587.5.

**Fig. 2:**
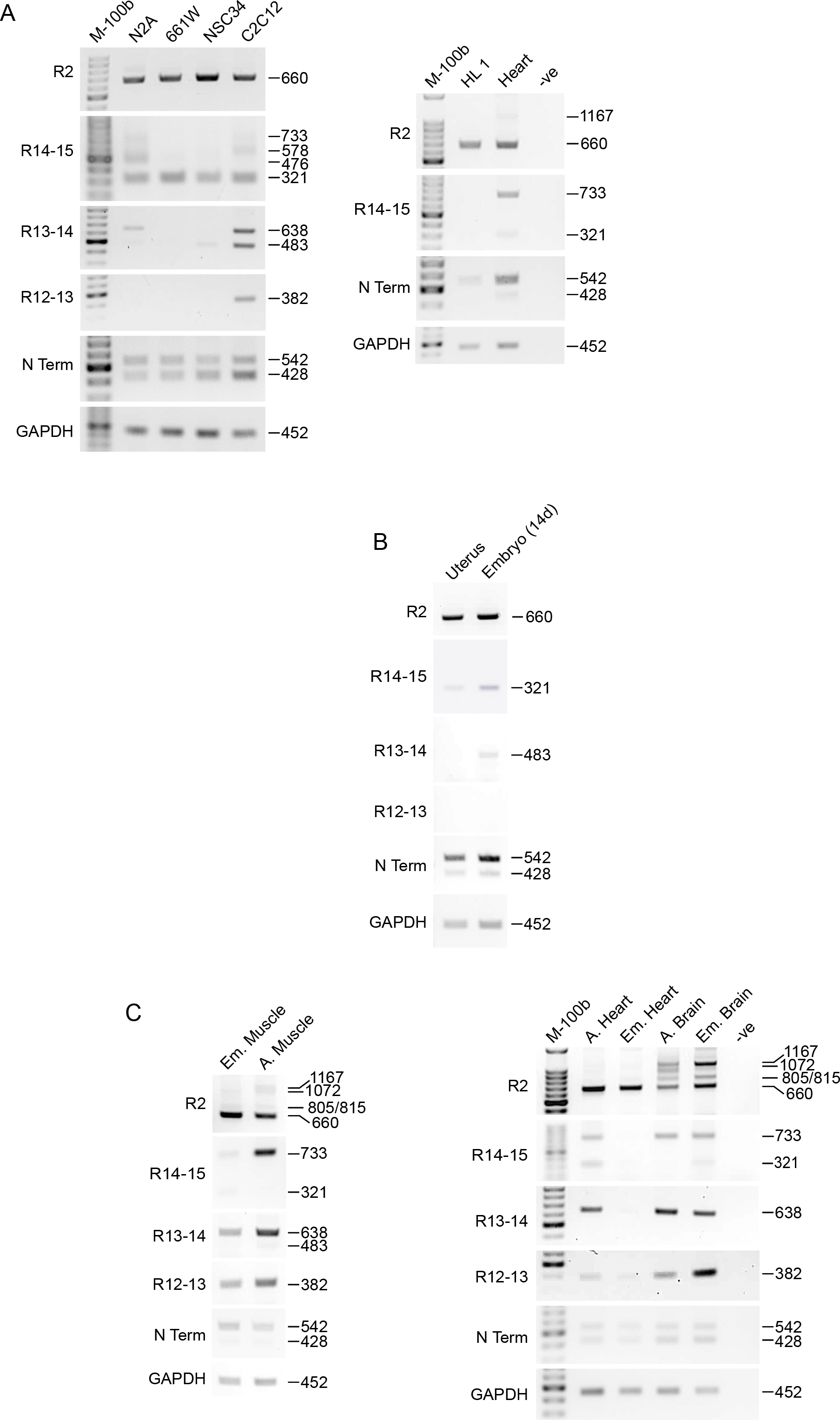
Expression analysis of RAPGEF1 isoforms in cell lines, and embryonic tissues. A) Three neuronal cell lines (N2A, 661W, NSC34), a myoblast line (C2C12), and cardiomyocyte line (HL-1), were examined. Adult heart was included along with HL-1 cell line for comparison. B) RAPGEF1 isoform expression in the 14d whole embryo and uterus. C) Comparison of isoform expression between embryonic and adult tissues, heart, skeletal muscle and brain.

### Differential expression of isoforms during development and differentiation

Examination of whole mouse embryo at 14 days, along with the uterus showed that they primarily express isoforms that have exon 3, and those that lack the cassette exons (isoform 3). Transcripts with exon 14 alone appear to be present at very low level (Fig. 2B). We had earlier found that isoform switching occurs during development of the mouse brain. Since some of the tissues in the adult, like heart, skeletal muscle, and brain showed isoforms with inclusion of cassette exons, we wished to examine their expression during development, by comparing tissues from 18d embryo with 8wk adult. Transcripts with exon 14 are totally absent in the embryonic heart, while adult heart shows their expression (Fig. 2C). While embryonic brain shows the presence of transcripts with exon 14 alone included, the adult brain shows expression of isoforms with inclusion of exons 12 & 13. Difference was also observed between embryonic and adult skeletal muscle, with adult tissue showing significantly higher levels of transcripts with cassette exons. Comparison of isoform expression in tissues of 1yr old mouse showed similar expression pattern to that seen in tissues of 8-week-old animals (Fig. S4)

These findings prompted us to examine isoform switching in an invitro differentiation model. C2C12 myoblasts can be differentiated to fuse and form myotubes in culture, and have been used as a good model to mimic skeletal muscle formation invivo [28]. RNA isolated from exponentially growing myocytes, and myotubes formed upon differentiation for 48 & 96 hrs were examined. Myocytes primarily express the isoform without cassette exons, and as they differentiate, expression of isoforms with incorporation of cassette exons increases with time (Fig. 3A). While the isoform with exon 14 alone decreases, those with exons 12 & 13 increase as observed in PCRs performed with exon specific primers. Expression of isoforms skipping exon 3 was not significantly altered during differentiation. Induction of differentiation was confirmed by increase in myogenin transcript levels. To determine if isoform switching was a feature of cell proliferation arrest, we examined isoform expression in cells arrested by serum starvation as well as density dependent arrest. Under both these situations, isoform having exon 14 alone, was expressed, but no expression was seen of isoform with exons 12,13 & 14, which increases upon differentiation. These results indicated that cassette exon incorporation is specific to the process of differentiation, and not simply cell proliferation arrest that accompanies differentiation.

**Fig. 3:**
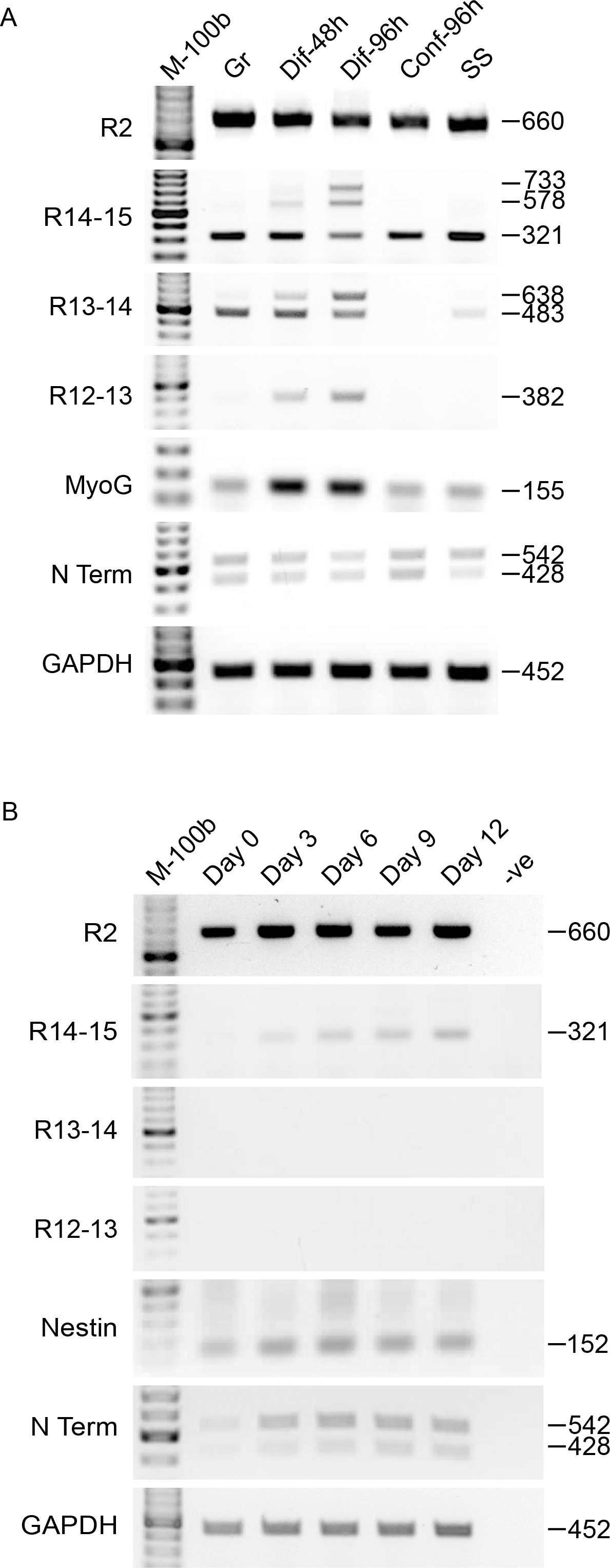
Isoform switching during myogenic differentiation of C2C12 cells and embryoid body (EB) formation of E14Tg2a, mouse ES cell line. A) Exponentially growing C2C12 cells, and those differentiated for 48, and 96 hrs were examined for expression of RAPGEF1 isoforms. Cells arrested by serum starvation and confluence were also compared. Myogenin (MyoG) expression was used to confirm differentiation. B) E14Tg2A cells were induced to form EBs for the indicated days and examined for RAPGEF1 isoform expression. Nestin expression was used to indicate ectoderm differentiation.

We also followed isoform expression during mESC differentiation. Growing E14-Tg2A cells primarily express only isoform 3 without any of the cassette exons (Fig.3B). But on inducing embryoid body (EB) formation for 12 days, when they begin to show lineage specific differentiation, we observed a gradual increase in the transcript with incorporation of exon 14. A corresponding increase was also observed in total RAPGEF1 levels, as seen by increase in products detected using the N-term primers. No expression was seen of isoforms with exons 12 & 13. Nestin levels were examined to indicate EB differentiation.

We compared our expression analysis with that obtained from publicly available RNA-seq data sets, and our own data on mESCs, which showed differential expression of predicted protein coding isoforms in mouse tissues (Table S1). Similar to that seen from our results, RAPGEF1 isoform lacking 12-14 cassette exons is ubiquitous, and was predominant in ESCs and embryonic tissues. Alternate isoforms showed similar trends in expression levels with highest levels of isoform 1 in the brain, isoform 2 in the skeletal muscle, and isoform 3 in the ovary, spleen and liver. Some variability in expression of the isoforms was seen compared to that in our analysis. This may be due to the short length of RNA-seq reads that hampers the discovery of contiguous exon connectivity for full-length alternatively spliced isoforms.

### Characterization of the amino acids incorporated by the cassette exons

To characterize the cassette exons which are incorporated during differentiation, we examined the amino acid sequence of the polypeptide inserted. A total of 170 amino acids are added with 52, 86, and 32 amino acids being contributed by exons 12, 13, and 14 respectively, which are highly conserved across species (Fig. S5). Analysis using NetSurfP-3.0 tool showed that most of this sequence forms a random coil, with high surface accessibility, and intrinsic disorder (Fig. 4A). Such polypeptides have been shown to form intracellular aggregates, or undergo phase separation [29, 30]. The 138 additional amino acids within the major brain isoform (isoform 1, containing exons 12 & 13) were cloned as a GFP fusion protein (GFP-138mBr) and expressed in HEK293T cells. While GFP is a soluble protein, localized in the nucleoplasm and cytoplasm, GFP-RAPGEF1 shows predominant cytoplasmic staining in HEK293T cells as described earlier (Fig. S6). The GFP-138mBr fusion protein localized throughout the cell, but formed distinct juxta-nuclear aggregates in the cytoplasm, suggestive of phase separation. Interestingly, it was observed that the sequence is extremely serine-rich (23%), (Fig. S5), while the average number of serines in proteins is around 7%. We therefore examined this sequence for the presence of predicted target sites of phosphorylation by serine kinases, and identified multiple sites with high probability of being phosphorylated (Fig. 4B). PKC, CKII, GSK3β, Cdk5, and DNA-PK are kinases predicted to phosphorylate these sites. We examined the fusion protein by SDS-PAGE, and western blotting to examine if it shows any indication of post-translational modification. In addition to a band of the expected 44 kDa, a band of about 50kDa was seen, when probed with GFP antibody. To test if the mobility shift was caused by phosphorylation, we observed its expression in the presence and absence of phosphatase inhibitors, OA, and PV. As shown in Fig. 4C, there was a significant increase in bands of retarded mobility upon inhibiting S/T phosphatases, but not Tyr phosphatases. It was therefore highly probable that this intrinsically disordered region in isoforms of RAPGEF1 expressed in select tissues is phosphorylated by S/T kinases.

**Fig. 4:**
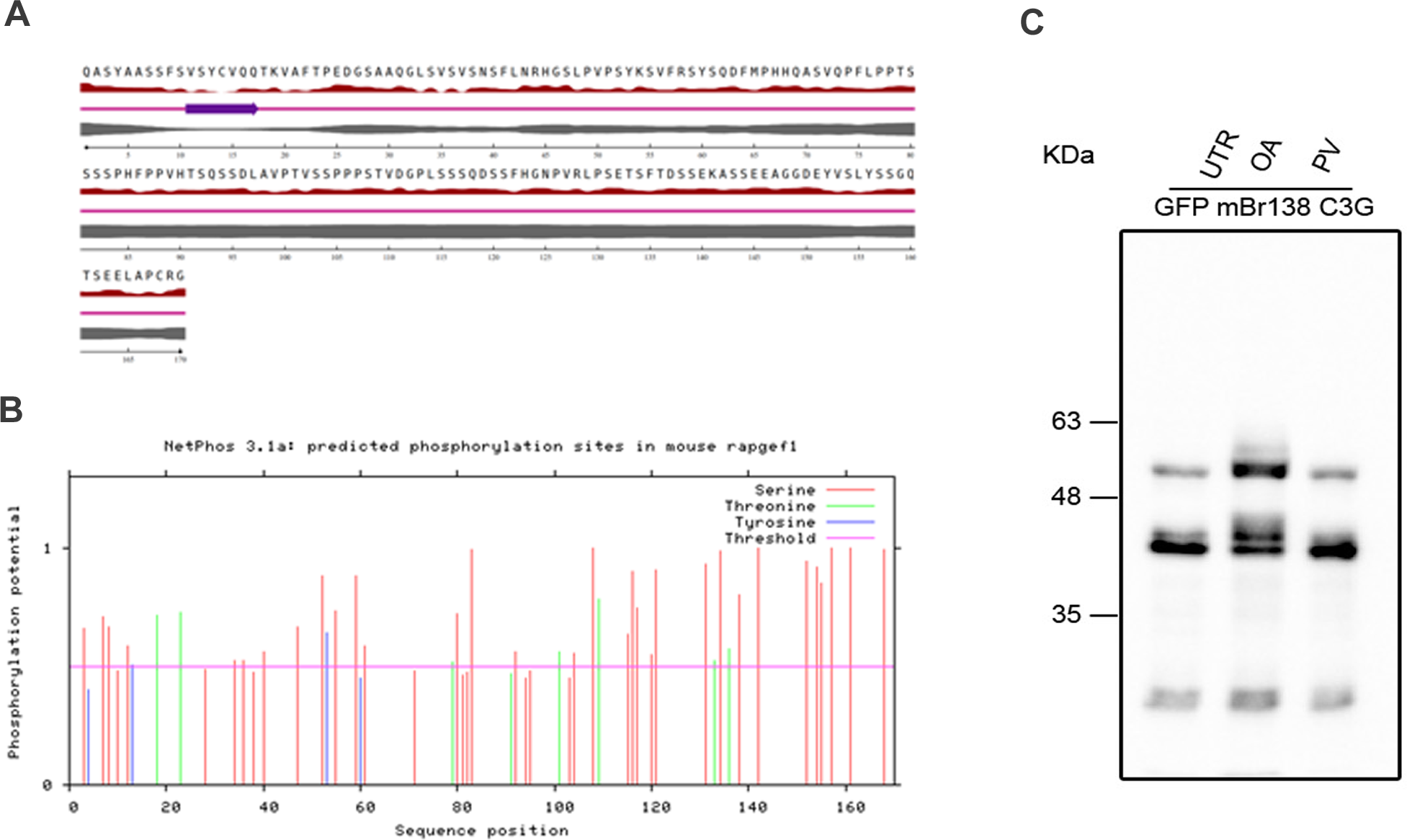
Peptide segment introduced by cassette exons is disordered, serine-rich, and highly phosphorylated. A) Prediction using NetSurfP - 3.0 - Protein secondary structure and relative solvent accessibility tool, indicates that this segment is intrinsically disordered. Lines from top to bottom indicate amino acid sequence, Relative surface accessibility, Secondary structure, Disorder, and residue numbering. B) Netphos prediction of target site phosphorylation in this disordered segment. C) Western blot showing mobility shift in a fusion protein of GFP with amino acids encoded by exons 12 & 13 cloned from mouse brain (GFP-138brC3G) upon treatment with Ser/Thr and Tyr phosphatase inhibitors, OA, okadaic acid & PV, pervanadate, respectively. UTR, untreated.

### The amino acids inserted by the cassette exons alter intramolecular interactions

The non-catalytic domains of RAPGEF1 are known to regulate access of the substrate and activity of the catalytic site through intramolecular interactions. Crk binding to the poly-proline tracts, and mutations that inhibit AIR from binding to the catalytic domain, can alter substrate binding, and catalytic activity [6, 10]. The 3D structure of full length RAPGEF1 has not been determined, and lack of structural information has obscured the functional consequences of its intramolecular and intermolecular interactions. The AlphaFold protein structure database provided a model structure for us to investigate intramolecular interactions within RAPGEF1 [23, 31]. Insilico analysis was therefore carried out to examine how the presence of the additional residues in isoforms 1 & 2 alter interaction of the AIR with the catalytic domain, compared to isoform 3, which does not have exons 12, 13 & 14. The amino acid sequence of the 3 isoforms used for our predictions are shown in Fig. S1. Residues within the AIR that are essential for interaction are within a short alpha helix, and are conserved across species. We examined this segment within the structure for possible formation of hydrophobic interactions with residues in the catalytic domain, and also specifically examined proximity of three invariant residues, M523, Y526, and M527, that have earlier been shown to be essential for inhibitory function of AIR [6]. It was seen that presence of segments contributed by cassette exons correlates with stronger interface between AIR and three alpha helices in the catalytic domain (CDα1, CDα2 and CD α3) (Fig.5, & Fig. S1). Hydrogen and hydrophobic bonds formed between the AIR and catalytic domain helices keep it in a closed confirmation, a mechanism that represses its GEF activity when unstimulated.

**Fig. 5:**
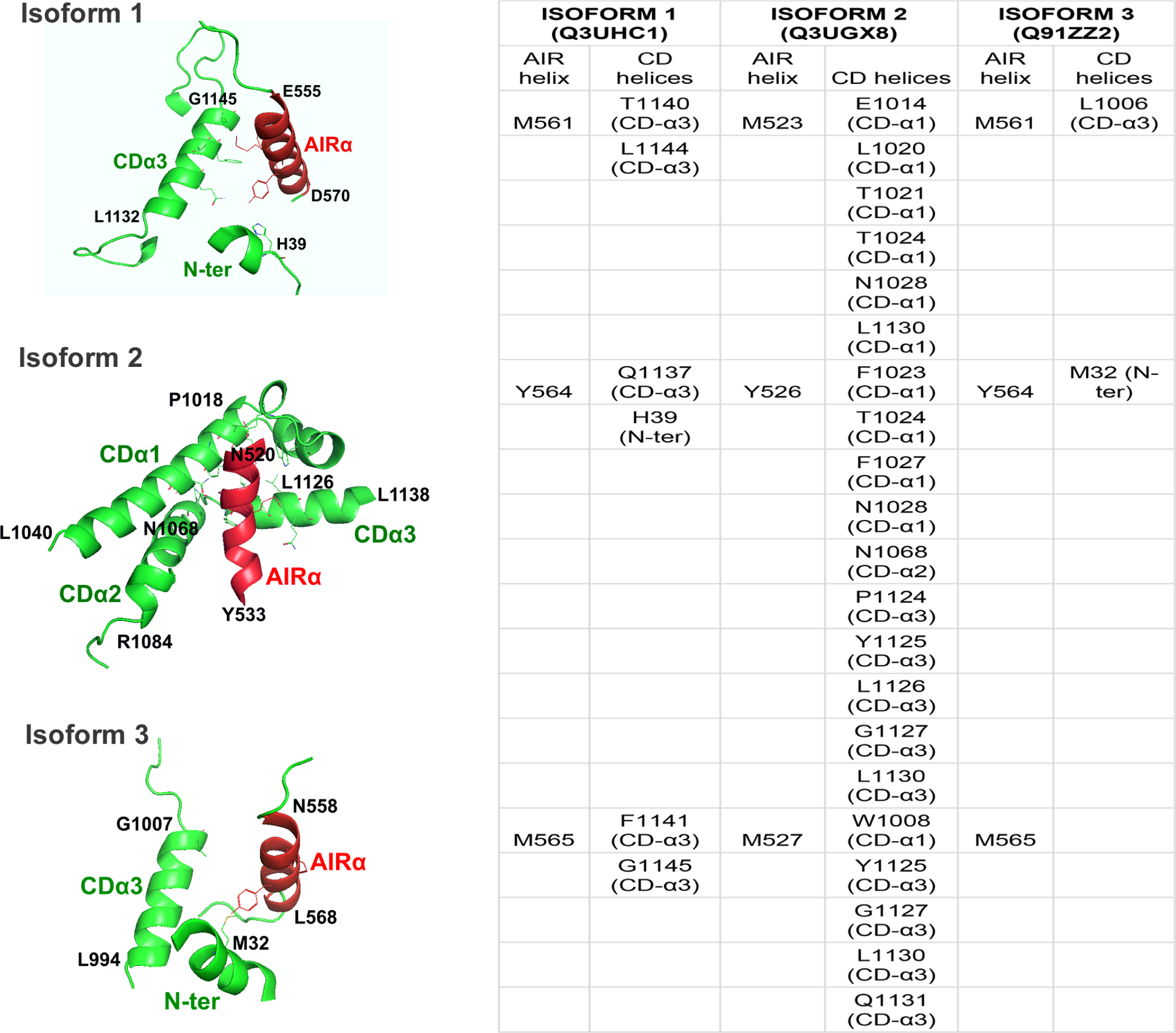
RAPGEF1 isoforms differ in intramolecular interaction between the AIR and their catalytic domain. AlphaFold derived 3D model structures of RAPGEF1 isoforms showing regions in its catalytic domain (green) that come into close proximity to its AIR helix (red). Predicted interactions of invariant AIR residues, M561, Y564, and M565 with those in the CD are shown. CDα 1,2, &3 indicate the alpha helices in the CD, which are identical in all the isoforms. (Refer Fig. S1 for sequence alignment and numbering, which differs between the isoforms due to alternate splicing). Amino acid numbers at the start and end of each helix are indicated as per numbering in the respective isoforms. Table lists all the amino acids in the catalytic domain with which the residues in the AIR helix (red) occur in close proximity (<5.0 Ang).

While, isoform 3 & 1 show interface with only 1 of the helices (CDα1), isoform 2 interfaces with all three helices. Studies suggest 6-8Ang distance being suggestive of interaction [32, 33]. Residues that are within the stringent cutoff of 5.0A distance, and likely to make hydrophobic interactions are shown in table (Fig. 5). The invariant residues within the AIR in isoform 2 are within 5.0 Ang distance from multiple residues in the catalytic domain, but in isoform 3, the helixes are displaced by over 20Ang, and therefore likely to enable easier access to the catalytic site. It was also noticed that a short helix in the beginning of the N-term, inserts in between the AIR and catalytic domain in the case of isoform 1, and 3, with the AIR possibly forming hydrophobic interactions with H39, and M32 respectively. Interestingly, it was observed that it was primarily the three invariant residues, in the AIR, that show likely interactions, as the number of contact sites did not differ when the whole helix was examined (data not shown). The helices in the catalytic domain we identified as coming in close contact with AIR helix are those where the structural predictions have been made with very high confidence limit (pLDDT score above 90%).

### Protein-protein interaction (PPI) analysis shows differences in interaction of RAP1 with RAPGEF1

GEFs are known to interact with target GTPases through their catalytic domains, and stabilize them in a nucleotide-free state, favouring replacement of GDP on the GTPases with GTP [34]. Structural analysis of RAPGEF1 in association with RAP1, a known target, has not been carried out, but mutational analysis has shown that RAP1 primarily interacts with its GEFs through amino acids in its SwI and SwII domains [35, 36]. Structures of nucleotide-free complexes of catalytic domains have been described for the RasGEF, SOS, and the RapGEF EPAC, showing that the GTPase-binding surface is entirely comprised within the-helical bowl-shaped Cdc25 homology domain(Cdc25HD), which is highly homologous to that of SOS [34, 37]. The REM domain forms a close-packed interaction with the Cdc25HD that is very similar in all known structures, whether this domain is bound or not to its cognate GTPase. Deletion analysis has shown that RAPGEF1 engages RAP1 through the Cdc25HD [3, 38]. The N-term region of RAPGEF1 is autoinhibitory, and therefore, we focused our analysis on the docking of isoforms 1,2 & 3 lacking N-term with RAP1A (residues 540 to 1086 in isoform 3, 540 to 1224 in isoform 1, and 502 to 1218 in isoform 2; shown in Fig. S1), from the structures of the full-length forms for ease of analysis. In the absence of experimentally determined structure of RAPGEF1 in complex with RAP1, we used the molecular docking tool, LZerD to analyze the possible effects of exons 12/13/14 inclusion on the interaction between RAPGEF1 isoforms and RAP1 protein. Using AlphaFold derived structures of RAPGEF1 and RAP1A, we subjected them to rigid body docking using LZerD pairwise protein–protein docking. We selected as binding sites all of the regions where the number of contacts with RAP1A for each residue was greater than 600. After calculating the distance between carbon α of the interacting residues, the best docked pose was detected according to the MM/GBSA server, with the free binding energy of −18.11 kcal/mol for RAPGEF1 isoform1, - 37.57 kcal/mol for RAPGEF1 isoform2 and −23.3 kcal/mol for RAPGEF1 isoform3. The docking results showed a certain selectivity in the interaction of RAPGEF1 with RAP1A. The results are shown in Fig. 6. Protein-protein interaction residues are defined as those which have a heavy atom closer than 5.0 Å to any atom in the docking partner [39], but the cutoff distance commonly used is 6 angstroms [32]. The rank sum scores for the isoforms 1,2 & 3 were 869, 446, and 398, respectively. Considering all residues between the two molecules that are within 5.4Ang, we identified differences in the predicted interaction of RAP1A with the various residues of the RAPGEF1 isoforms. Interestingly, we observed closer and more interactions of RAP1A through amino acids in its switch domains with residues of isoform 3, which was predicted to be in a more open confirmation relative to isoform. In this case, the contact sites fall within the regions identified earlier in RAP1. The C-term tail that is modified for membrane localization is not involved in interactions. RAP1 primarily interacts with an a-helix in RAPGEF1 formed by residues 788 to 826. This region encompasses the NES and GID identified by us earlier [9, 15].

**Fig. 6:**
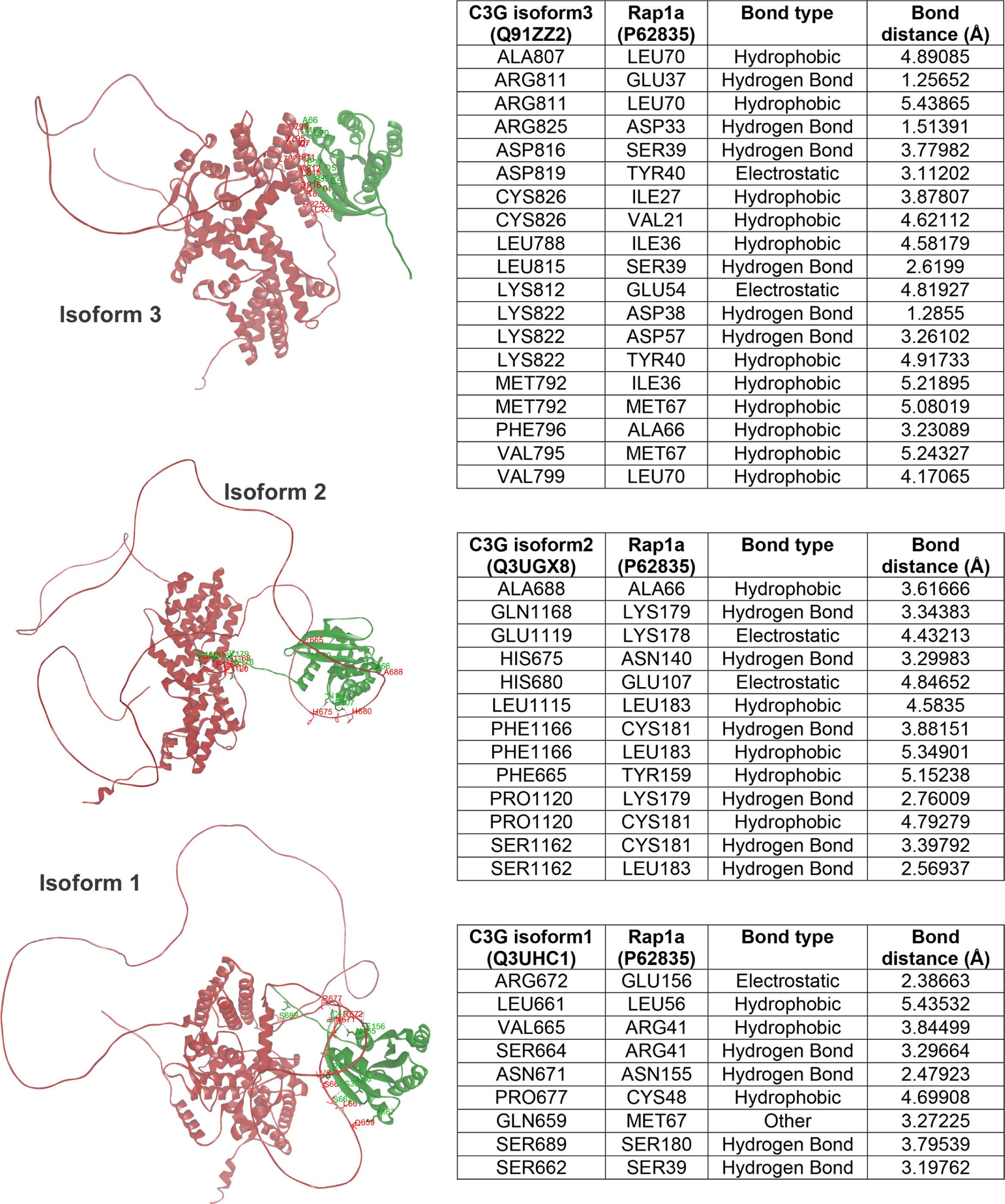
RAP1 docks differentially to RAPGEF1 isoforms with and without cassette exons. Images show best fit docking results of RAP1(green) with RAPGEF1 (red), obtained from the LZerD protein docking suite. Only the C-term half of RAPGEF1 isoforms were used for the PPI as mentioned in the methods. 5.4Ang was used as maximum distance between amino acid pairs that dock in close proximity, and the nature of their interactions is listed in the table.

Interestingly, we observed that the RAP1 docking sites are different in the case of isoform 1, and 2, which have additional residues, that form an IDR. In case of isoform 2, all the interacting residues in RAP1 are outside the switch regions. They primarily interact with two distinct regions, one of them being the IDR formed by the cassette exons, and the other with regions in the CD. In case of isoform 1, all interactions are restricted to the IDR of the cassette exons, without interaction with the CD. Our PPI analysis therefore showed interaction of residues contributed by cassette exons in isoforms 1, and 2 with RAP1, altering its ability to orient itself with catalytic surface of RAPGEF1. The C-term tail of RAP1 also makes contacts with residues in these isoforms, indicating that it may not be available for its other functions like membrane anchoring.

## Discussion

Evaluation of the expression of tissue-specific isoforms of genes known to show ubiquitous expression, enables greater insight into functions served by distinct isoforms in specialized cell types as alternatively spliced isoforms exhibit different intra, and inter molecular interactions [40]. RNA sequence profiling of mouse and human tissues has shown the expression of five protein coding isoforms of Rapgef1, but its tissue specific expression has not been systematically investigated. We examined expression patterns of isoforms across many mouse tissues and cell types by carrying out semi-quantitative RT-PCR that can detect distinct alternately spliced isoforms. Of the 27 exons in the gene, alternate splicing of exons occurs at two locations, i.e. exon 3, and exons 12,13, & 14. Splicing of exon 3 is universal, with all tissues showing transcripts with and without this exon, though their relative levels vary. Amino acids encoded by exon 3 are well conserved across species, and is predicted to form a largely unstructured region, with a short helix. How residues in this exon contribute to RAPGEF1 activity, and signaling functions requires further investigation. The transcripts expressed in some adult tissues like brain, testis, heart and skeletal muscle have one or more cassette exons (12, 13 &14), and their insertion does not alter reading frame, resulting in the presence of additional amino acids (of varying number) following the last poly-proline tract adjacent to the AIR. The presently available data bases (NCBI & Ensemble), predict generation of isoforms with multiple combinations of the three cassette exons. We demonstrate tissue specific expression of some of the predicted isoforms, and also provide experimental evidence for expression of two novel and unpredicted transcripts, one with incorporation of cassette exon 14 alone, and the other with cassette exons 12 and 14, but missing exon 13. While the former is present at low levels in tissues with primary expression of the canonical isoform lacking cassette exons, the latter is expressed in neuroblastoma cell line, N2A. Splicing at the two specific hotspots is conserved in human and rat RAPGEF1 [21, 22]. Western blots have generally shown a major band at 140kDa, and slow-moving polypeptides, or a smear, which could correspond either to the longer isoforms, or post translationally modified protein [7, 15]. Polypeptides of 175kDa, corresponding to isoform 1, were expressed in adult mouse brain, and documented by sequencing [22]. Antibodies that are presently available have been generated using domains common to all isoforms, and therefore cannot distinguish between them, unless they differ significantly in size in western blots. Therefore, our study could not be supplemented by showing expression of polypeptides derived from the variant isoforms in all the tissues.

Our results indicate that isoform switch is associated with differentiation, and not cell cycle arrest, as we did not observe significant expression of the cassette exon containing transcripts in cells arrested by serum starvation or contact inhibition of C2C12 cells. There are other examples where a ubiquitously expressed protein regulates muscle differentiation through specific splicing [41]. Active RAP1A protein acts as a potent inhibitor of the differentiation program [42]. Isoform switching towards more inactive forms during myogenic differentiation is therefore likely to regulate RAP1 activity to aid differentiation. Our work on mESCs showed that is essential for maintaining Erk activity, and Erk inhibition is known to be required for promoting myocyte fusion [43]. Switching isoform expression may therefore be an additional mechanism to regulate signaling to enable differentiation.

It is known that amino acids incorporated due to inclusion of cassette exons in a tissue specific manner generally form disordered segments that embed binding motifs which can alter interactions with other proteins [44]. The amino acids encoded by the cassette exons of RAPGEF1 also form a disordered segment. Our results show that these tissue specific segments are modified by reversible phosphorylation at multiple serines, which may further contribute to intramolecular and intermolecular interactions. Phosphorylation of serine-rich IDRs is known to alter activity of proteins [45, 46]. Cdk5, a kinase involved in neuronal differentiation phosphorylates RAPGEF1 to alter RAP1 activity [47]. We identified Cdk5 target sites in the insert of the major isoform expressed in adult brain, suggesting that RAPGEF1 activity is regulated in mature neurons through isoform switching and phosphorylation.

Intrinsically disordered regions in proteins promote phase separation to enable compartmentalization, a property important in multiple biological processes. They are common among proteins involved in transcription, RNA processing, and chromatin organization [48, 49]. We observed that the amino acids introduced by the cassette exons of RAPGEF1 have the ability to form granular, membrane-less structures when fused to GFP, suggesting that they contribute to its specific compartmentalization. Our earlier work demonstrated the association of RAPGEF1 with chromatin and with nuclear speckles [15, 50], suggesting that the variant isoforms may differ in some of these properties and their related intracellular functions.

The auto-inhibitory region of RAPGEF1 folds over and binds to residues in the catalytic domain, keeping its activity in check [6]. The additional residues in the longer isoforms, which are inserted following the last poly-proline tract, and adjacent to the AIR, may alter binding of AIR with the catalytic domain, or alter ability of SH3 containing proteins from binding to the adjacent poly-proline tracts. Our analysis of the predicted structure of RAPGEF1 from AlphaFold, indicated that isoforms with longer segments between the CBR and catalytic domain, enabled better interaction of the AIR (residues K521 to L530, numbering as in isoform 202) with the catalytic domain, and therefore may maintain a less active confirmation. This insilico analysis is further validated by the fact that the same AIR helix identified through mutagenesis [6] comes in close proximity with helices in the catalytic domain. Other studies have also shown the utility of AlphaFold to identify structures of IDR containing proteins [51, 52].

Intramolecular interaction between the N-term segment (aas 4-245) and the catalytic domain, contribute to increase in catalytic activity of RAPGEF1, but the exact residues involved were not identified [6]. We observed that in the predicted structure of RAPGEF1, residues in the N-term form a helix which comes in close proximity to the AIR and catalytic helix, in isoform 1 and isoform 3, but not in case of isoform 2. The presence of cassette exons not only alters AIR-catalytic domain interaction, but also affects the positioning of the N-term, which also contributes to regulation of catalytic activity. Carabias et al showed that the N-term interacts specifically with the REM domain, but we find that in addition, a short helix in the N-term comes in close proximity, and could interact with AIR residues important for auto-inhibition. It is suggestive of forming a wedge between the AIR and the catalytic domain specifically in isoforms 1 and 3. Residues in the REM region identified for interaction with the NTD, are E740 & E794 (in isoform 3) of mouse. E794 is within the region identified by us earlier as the nuclear export sequence and also GSK3β interaction domain [9, 15]. It appears that residues in the REM domain serve multiple functions, and its interaction with NTD may regulate nuclear export, as well as interaction with other molecules like GSK3β. Whether the N-term domain regulates these properties of RAPGEF1 is subject to further experimental investigation. Our findings provide evidence for multiple intramolecular interactions within domains of RAPGEF1 that could regulate its activity levels as well as intermolecular interactions in a graded fashion. From our docking studies, it appears that residues included due to incorporation of cassette exons also serve to interact with RAP1, and orient it unfavourably for catalytic activity. It is also seen that the extreme C-term of RAP1, which has a poly-basic region, and is modified for cell membrane anchoring, docks with residues in the longer isoform of RAPGEF1.

Structures of several nucleotide-free GTPase-GEF complexes have been described [53], but that of RAPGEF1 complexed with Rap is not known. Several GEF catalytic domains have been crystallized both alone and in complex with their target GTPase, showing that they barely undergo any conformational changes on their own. Regulation is mostly through intramolecular and intermolecular interactions. Non-catalytic domains can fold over and prevent access to the GTP binding site, and this can happen in a graded manner for achieving different levels of catalytic activity. The linker region introduced by the additional exons in RAPGEF1, rather than enabling more flexibility, appears to aid an autoinhibited conformation, as well as altering Rap 1 binding at the catalytic site. High conservation of the amino acids incorporated by the cassette exons supports their essential role in some of the adult tissues. The 14d mouse embryo, 18d embryonic brain and skeletal muscle as well as mouse ESCs predominantly express isoform 203, and isoform switching occurs in some adult tissues like the brain, testis and skeletal muscle, but not in others like kidney, thymus, liver and lung. Development of isoform specific antibodies, and additional studies on isoform specific knockout mice will reveal tissue specific functions of these variant isoforms.

## Conclusion

Our results show that splice variants of RAPGEF1 are differentially expressed during development in a tissue specific manner. All embryonic tissues, and some adult tissues like lung, liver, spleen, and kidney showed predominant expression of the canonical isoform, and highly differentiated tissues with few proliferating cells expressed the longer isoforms containing cassette exons 12-14, suggesting that isoform switching is important for differentiation and maintenance of some adult tissue types. Our analysis of AlphaFold models of various isoforms provided structural evidence for differences in intramolecular interactions, possibly regulating RAPGEF1 catalytic activity and substrate binding. Our results indicate a novel means of regulating RAPGEF1 function, through addition of different combinations of cassette exons. and yield insights into understanding differences in the properties of alternate isoforms.

## Acknowledgements

Thank Vishnu Vijay for gift of Tg2A EB samples. We thank Dr. Tej Sowpati for providing help with the bioinformatics. VR acknowledges CSIR for award of ES scheme, 21(1089)/19/EMR-II.

## Author contributions

AV, AG, NK, and GG performed the experiments, analyzed the data, and prepared the figures. VR conceived & supervised the study, analyzed the data, and prepared the manuscript. All authors read and approved the manuscript.

## Abbreviations

AIR: Auto-inhibitory region
GEF: guanine nucleotide exchange factor
CBR: CRK binding region
OA: okadaic acid
AS: alternate splicing
REM: Ras exchanger motif
P-P: poly-proline
CD: catalytic domain
CDC25HD: CDC25 homology domain
PPI: protein-protein interaction
IDR: intrinsically disordered region

