## Supplemental figs and table for "Development and tissue specific expression of RAPGEF1 (C3G) transcripts having exons encoding disordered segments with predicted regulatory function"

```
                                                1|2
NP_001034175.1_Isoform2  MSSGLGLRRSPEMSGKIEKADSQRSHLSSFTMKLMDKFHSPKIKRTPSKKGKPAEVSKIP 60
NP_001034176.1_Isoform1  MSSGLGLRRSPEMSGKIEKADSQRSHLSSFTMKLMDKFHSPKIKRTPSKKGKPAEVSKIP 60
NP_473391.1_Isoform3     MSSGLGLRRSPEMSGKIEKADSQRSHLSSFTMKLMDKFHSPKIKRTPSKKGKPAEVSKIP 60
                          2|3                                      3|4
NP_001034175.1_Isoform2  EKPVS--------------------------------------KNLCWLEEKEKEVVSAL 82
NP_001034176.1_Isoform1  EKPVSKEARDRFLPEGYPIPLDLEQQAVEFMSTSAVASRSQRQKNLCWLEEKEKEVVSAL 120
NP_473391.1_Isoform3     EKPVSKEARDRFLPEGYPIPLDLEQQAVEFMSTSAVASRSQRQKNLCWLEEKEKEVVSAL 120
                                                                4|5
NP_001034175.1_Isoform2  RYFKTIVDKMAIDKKVLEMLPGSASKVLEAILPLVQTDPRIQHSSALSSCYSRVYQSLAN 142
NP_001034176.1_Isoform1  RYFKTIVDKMAIDKKVLEMLPGSASKVLEAILPLVQTDPRIQHSSALSSCYSRVYQSLAN 180
NP_473391.1_Isoform3     RYFKTIVDKMAIDKKVLEMLPGSASKVLEAILPLVQTDPRIQHSSALSSCYSRVYQSLAN 180
                                                            5|6
NP_001034175.1_Isoform2  LIRWSDQVMLEGVNSEDKEMVTTVKGVIKAVLDGVKELVRLTIEKQGRPSPTSPVKPSSP 202
NP_001034176.1_Isoform1  LIRWSDQVMLEGVNSEDKEMVTTVKGVIKAVLDGVKELVRLTIEKQGRPSPTSPVKPSSP 240
NP_473391.1_Isoform3     LIRWSDQVMLEGVNSEDKEMVTTVKGVIKAVLDGVKELVRLTIEKQGRPSPTSPVKPSSP 240
                             6|7                                               7|8
NP_001034175.1_Isoform2  ASKPDGQPELPLTDREMEILNKTTSVSPSAELLPDSTSEEVAPPKPPLPGIRVVDNSPPA 262
NP_001034176.1_Isoform1  ASKPDGQPELPLTDREMEILNKTTSVSPSAELLPDSTSEEVAPPKPPLPGIRVVDNSPPA 300
NP_473391.1_Isoform3     ASKPDGQPELPLTDREMEILNKTTSVSPSAELLPDSTSEEVAPPKPPLPGIRVVDNSPPA 300
                                                                8|9
NP_001034175.1_Isoform2  LPPKKRQSAPSPTRVAVVAPMSRATSGSSLPVGINRQDFDVECYTQRRLSGGSRSCGGES 322
NP_001034176.1_Isoform1  LPPKKRQSAPSPTRVAVVAPMSRATSGSSLPVGINRQDFDVECYTQRRLSGGSRSCGGES 360
NP_473391.1_Isoform3     LPPKKRQSAPSPTRVAVVAPMSRATSGSSLPVGINRQDFDVECYTQRRLSGGSRSCGGES 360
                                                                 9|10
NP_001034175.1_Isoform2  PRLSPCSSTGKLSRSDEQLSSLDRDSGQCSRNTSCETLDHYDPDYEFLQQDLSNADQIPP 382
NP_001034176.1_Isoform1  PRLSPCSSTGKLSRSDEQLSSLDRDSGQCSRNTSCETLDHYDPDYEFLQQDLSNADQIPP 420
NP_473391.1_Isoform3     PRLSPCSSTGKLSRSDEQLSSLDRDSGQCSRNTSCETLDHYDPDYEFLQQDLSNADQIPP 420

NP_001034175.1_Isoform2  QAACNLSPLPESLGESGPPFLGHPFQLPLGSCLQQEGQQTDTPPALPEKKRRSAVSQTTD 442
NP_001034176.1_Isoform1  QAACNLSPLPESLGESGPPFLGHPFQLPLGSCLQQEGQQTDTPPALPEKKRRSAVSQTTD 480
NP_473391.1_Isoform3     QAACNLSPLPESLGESGPPFLGHPFQLPLGSCLQQEGQQTDTPPALPEKKRRSAVSQTTD 480

NP_001034175.1_Isoform2  SSGCRVSYERHPSQYDNISEGDLQNPVPVQPVPYPPFAAVLPFQQGASSASAEFVGDFSV 502
NP_001034176.1_Isoform1  SSGCRVSYERHPSQYDNISEGDLQNPVPVQPVPYPPFAAVLPFQQGASSASAEFVGDFSV 540
NP_473391.1_Isoform3     SSGCRVSYERHPSQYDNISEGDLQNPVPVQPVPYPPFAAVLPFQQGASSASAEFVGDFSV 540
                                           10|11
NP_001034175.1_Isoform2  PELAGDTEKPPPLPEKKNKHMLAYMQLLEDYSEPQPSMFYQTPQSEHIYQQKNKMLMEVY 562
NP_001034176.1_Isoform1  PELAGDTEKPPPLPEKKNKHMLAYMQLLEDYSEPQPSMFYQTPQSEHIYQQKNKMLMEVY 600
NP_473391.1_Isoform3     PELAGDTEKPPPLPEKKNKHMLAYMQLLEDYSEPQPSMFYQTPQSEHIYQQKNKMLMEVY 600
                                                    11|12
NP_001034175.1_Isoform2  GFSESFCGSDSTQELAPPPALPPKQRQLQASYAASSFSVSYCVQQTKVAFTPEDGSAAQG 622
NP_001034176.1_Isoform1  GFSESFCGSDSTQELAPPPALPPKQRQLQASYAASSFSVSYCVQQTKVAFTPEDGSAAQG 660
NP_473391.1_Isoform3     GFSESFCGSDSTQELAPPPALPPKQRQL-------------------------------- 628
                                             12|13
NP_001034175.1_Isoform2  LSVSVSNSFLNRHGSLPVPSYKSVFRSYSQDFMPHHQASVQPFLPPTSSSSPHFPPVHTS 682
NP_001034176.1_Isoform1  LSVSVSNSFLNRHGSLPVPSYKSVFRSYSQDFMPHHQASVQPFLPPTSSSSPHFPPVHTS 720
NP_473391.1_Isoform3     ------------------------------------------------------------ 628
                                                                        13|14
NP_001034175.1_Isoform2  QSSDLAVPTVSSPPPSTVDGPLSSSQDSSFHGNPVRLPSETSFTDSSEKASSEEAGGDEY 742
NP_001034176.1_Isoform1  QSSDLAVPTVSSPPPSTVDGPLSSSQDSSFHGNPVRLPSETSFTDS-------------- 766
NP_473391.1_Isoform3     ------------------------------------------------------------ 628
                                        14|15                         15|16
NP_001034175.1_Isoform2  VSLYSSGQTSEELAPCRGEPPSGKDGHPRDPSVSSASGKDSRENGERSPKSLDGLESAQS 802
NP_001034176.1_Isoform1  ------------------EPPSGKDGHPRDPSVSSASGKDSRENGERSPKSLDGLESAQS 808
NP_473391.1_Isoform3     ------------------EPPSGKDGHPRDPSVSSASGKDSRENGERSPKSLDGLESAQS 670
                                                 16|17                        17|18
NP_001034175.1_Isoform2  EEEVDELSLIDHNEIMARLTLKQEGDDGPDVRGGSGDILLVHATETDRKDLVLYCEAFLT 862
NP_001034176.1_Isoform1  EEEVDELSLIDHNEIMARLTLKQEGDDGPDVRGGSGDILLVHATETDRKDLVLYCEAFLT 868
NP_473391.1_Isoform3     EEEVDELSLIDHNEIMARLTLKQEGDDGPDVRGGSGDILLVHATETDRKDLVLYCEAFLT 730
                                        18|19                            19|20
NP_001034175.1_Isoform2  TYRTFISPEELIKKLQYRYEKFSPFADTFKKRVSKNTFFVLVRVVDELCLVELTEEILKL 922
NP_001034176.1_Isoform1  TYRTFISPEELIKKLQYRYEKFSPFADTFKKRVSKNTFFVLVRVVDELCLVELTEEILKL 928
NP_473391.1_Isoform3     TYRTFISPEELIKKLQYRYEKFSPFADTFKKRVSKNTFFVLVRVVDELCLVELTEEILKL 790
                                                                    20|21
NP_001034175.1_Isoform2  LMELVFRLVCSGELSLARVLRKNILDKVDQKKLLRCAHSDQPLAARGVAARPGTLHDFHS 982
NP_001034176.1_Isoform1  LMELVFRLVCSGELSLARVLRKNILDKVDQKKLLRCAHSDQPLAARGVAARPGTLHDFHS 988
NP_473391.1_Isoform3     LMELVFRLVCSGELSLARVLRKNILDKVDQKKLLRCAHSDQPLAARGVAARPGTLHDFHS 850
                                         21|22                                22|23
NP_001034175.1_Isoform2  HEIAEQLTLLDAELFYKIEIPEVLLWAKEQNEEKSPNLTQFTEHFNNMSYWVRSIIMLQE 1042
NP_001034176.1_Isoform1  HEIAEQLTLLDAELFYKIEIPEVLLWAKEQNEEKSPNLTQFTEHFNNMSYWVRSIIMLQE 1048
NP_473391.1_Isoform3     HEIAEQLTLLDAELFYKIEIPEVLLWAKEQNEEKSPNLTQFTEHFNNMSYWVRSIIMLQE 910
                                      23|24                                   24|25
NP_001034175.1_Isoform2  KAQDRERLLLKFIKIMKHLRKLNNFNSYLAILSALDSAPIRRLEWQRQTSEGLAEYCTLI 1102
NP_001034176.1_Isoform1  KAQDRERLLLKFIKIMKHLRKLNNFNSYLAILSALDSAPIRRLEWQRQTSEGLAEYCTLI 1108
NP_473391.1_Isoform3     KAQDRERLLLKFIKIMKHLRKLNNFNSYLAILSALDSAPIRRLEWQRQTSEGLAEYCTLI 970
                                                  25|26
NP_001034175.1_Isoform2  DSSSSFRAYRAALSEVEPPCIPYLGLILQDLTFVHLGNPDYIDGKVNFSKRWQQFNILDS 1162
NP_001034176.1_Isoform1  DSSSSFRAYRAALSEVEPPCIPYLGLILQDLTFVHLGNPDYIDGKVNFSKRWQQFNILDS 1168
NP_473391.1_Isoform3     DSSSSFRAYRAALSEVEPPCIPYLGLILQDLTFVHLGNPDYIDGKVNFSKRWQQFNILDS 1030
                                  26|27
NP_001034175.1_Isoform2  MRCFQQAHYEIRRNDDIINFFNDFSDHLAEEALWELSLKIKPRNITRRKTDREEKT 1218
NP_001034176.1_Isoform1  MRCFQQAHYEIRRNDDIINFFNDFSDHLAEEALWELSLKIKPRNITRRKTDREEKT 1224
NP_473391.1_Isoform3     MRCFQQAHYEIRRNDDIINFFNDFSDHLAEEALWELSLKIKPRNITRRKTDREEKT 1086
```

Fig. S1: Amino acid sequence and alignment of residues in the three RAPGEF1 isoforms: Alignment of residues derived from the 27 exons of mouse gene in the three experimentally verified isoforms. The isoform IDs given are from NCBI. Isoform 2 lacks exon 3, but has exons 12,13, & 14. Isoform 1 has exon 3, but lacks exon 14. Isoform 3 has exon 3, but lacks the cassette exons, 12, 13 & 14.The N-term helix involved in interaction with the AIR, and has a positive regulatory role, is highlighted in blue, and the P_P tracts are marked in yellow. The AIR is highlighted in orange, the REM in light green and the CDC25 HD in dark green. Amino acids from the cassette exons, 12, 13 & 14, are shown in red. The NES/G1D is indicated in bold case. Residues shown within the red dotted boxes in the CDC25HD, are the three alpha helices that are involved in interaction with the AIR helix. The arrowhead indicates the tyrosine which undergoes regulatory phosphorylation.

Fig. S1

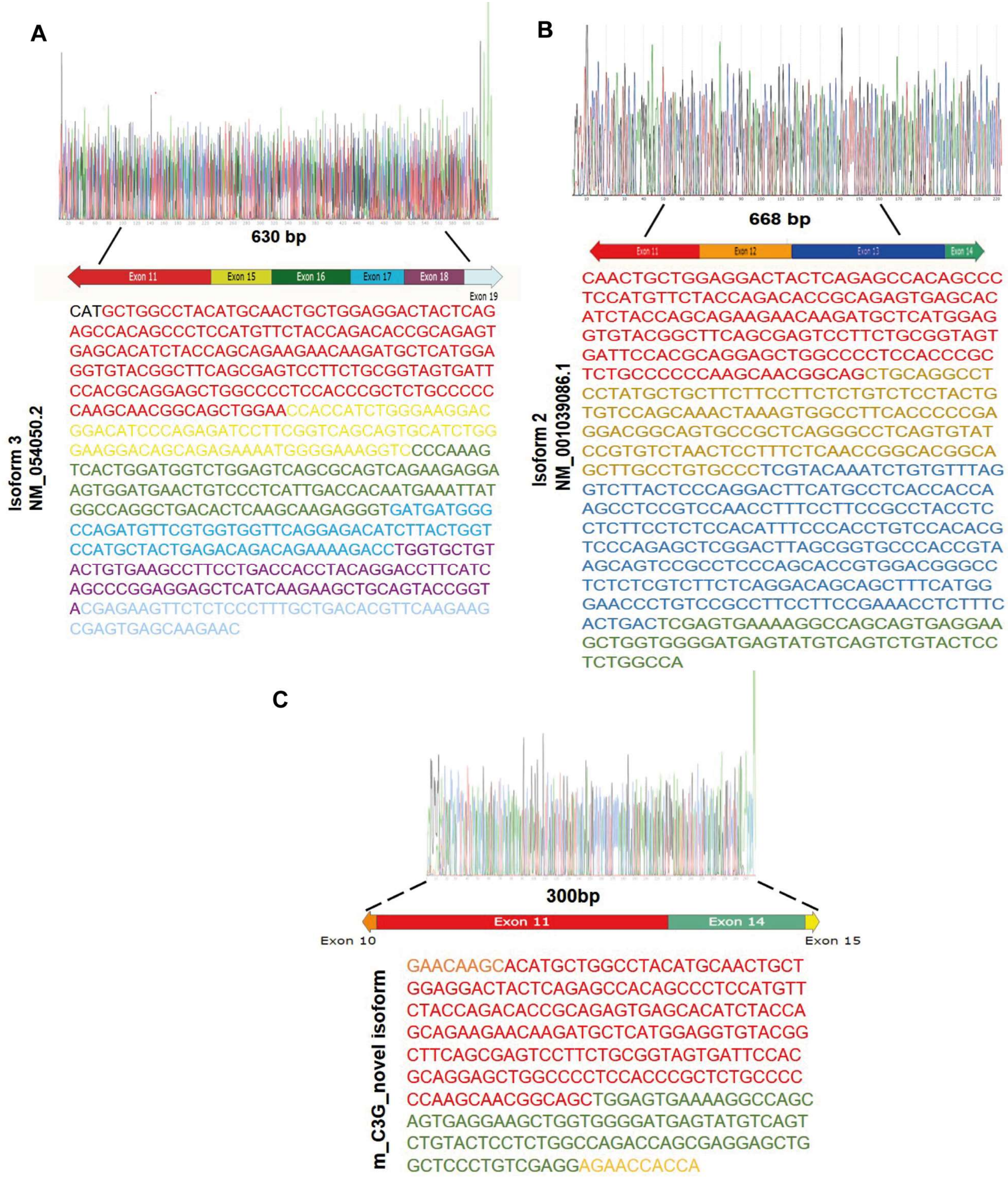


**Fig. S2 Sequencing results confirming that the PCR amplicons arise from various C3G transcripts: Fragments exciced from the gel after amplification were sequenced to confirm alternate splicing of the cassette exons. A. The 660bp amplicon obtained from ovary using forward (F), and reverse primer,(R2) had exon 15 (yellow sequence) contiguous with exon 11(red) confirming isoforms lacking cassette exons (C3G isoform 3.) B. Chromatogram showing sequence of the 733bp product obtaing from the brain using primers F, and reverse 14/15. This amplicon shows the presence of all 3 cassette exons, 12, 13 & 14 (isoform 2). C. Sequence of the 321bp product obtained from using F, and R14/15 primers shows exon 14 spliced to exon 11. This is a novel variant not predicted earlier.**

**Fig S2**

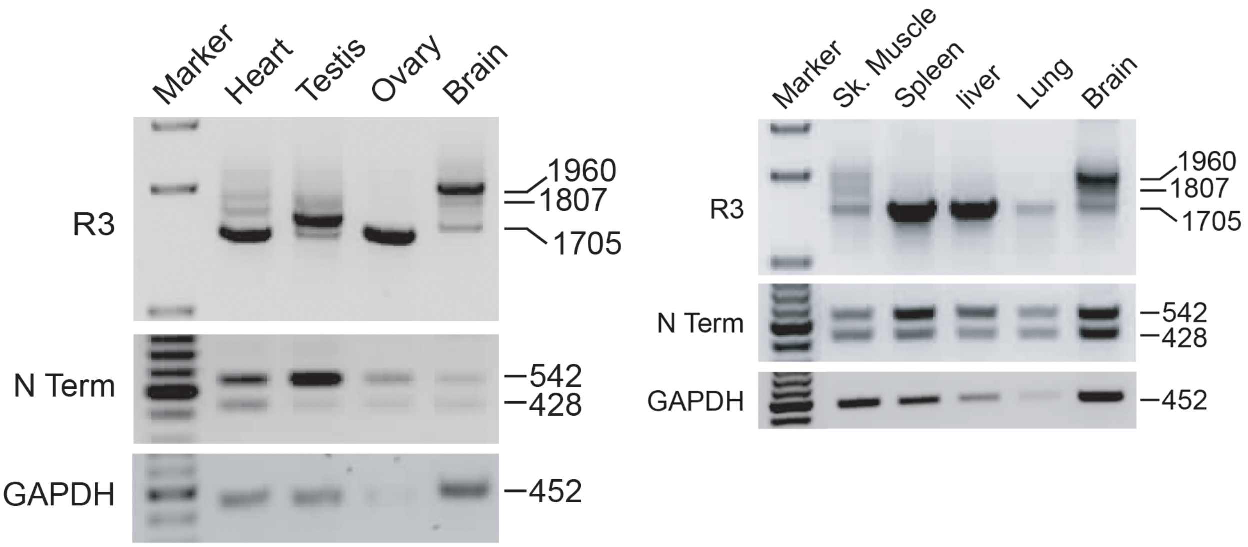


Fig.S3. Expression of RAPGEF1 isoforms in various tissues identified using an alternate reverse primer (R3). Tissues from adult mouse differ in expression of various isoforms including or excluding exons 12, 13, & 14, detected using an alternate reverse primer, R3 located in the last exon (Ref. Fig 1C). Tissues differ in expression of amplicons of various lengths.

Fig. S3

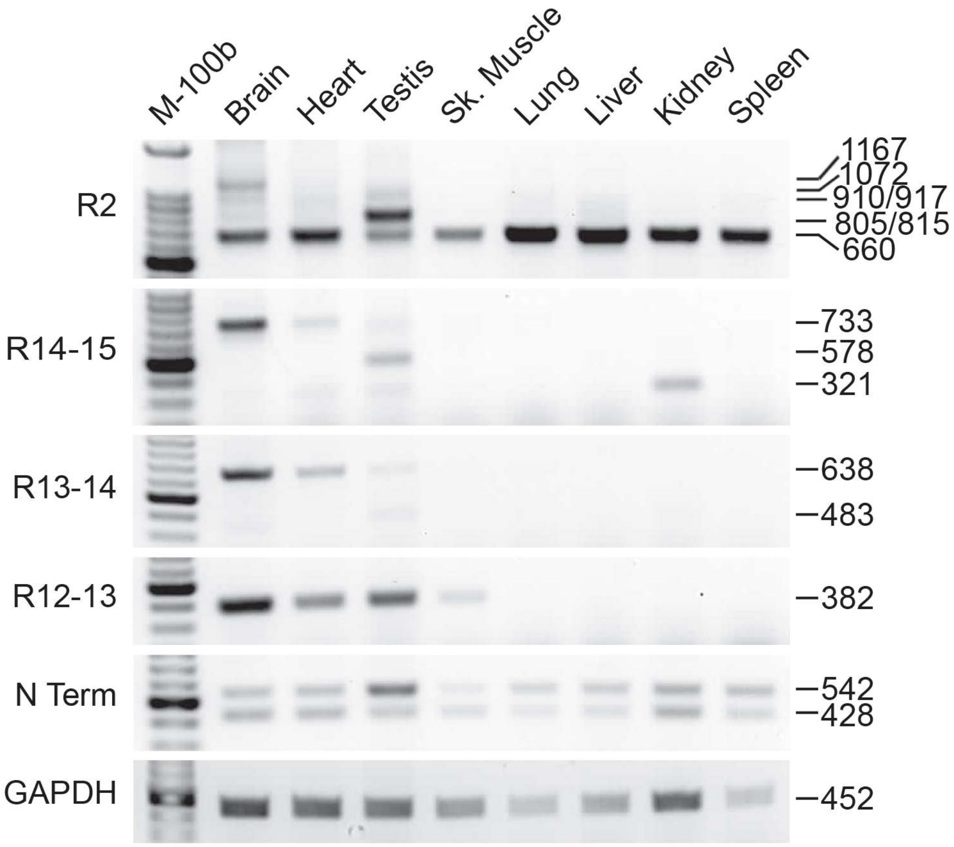


Fig. S4. Tissue specific expression of RAPGEF1 isoforms in one-year old mouse. Indicated primers were used to identify expression of splice-site variants in various tissues. Sizes of amplicons are indicated on the right.

Fig. S4

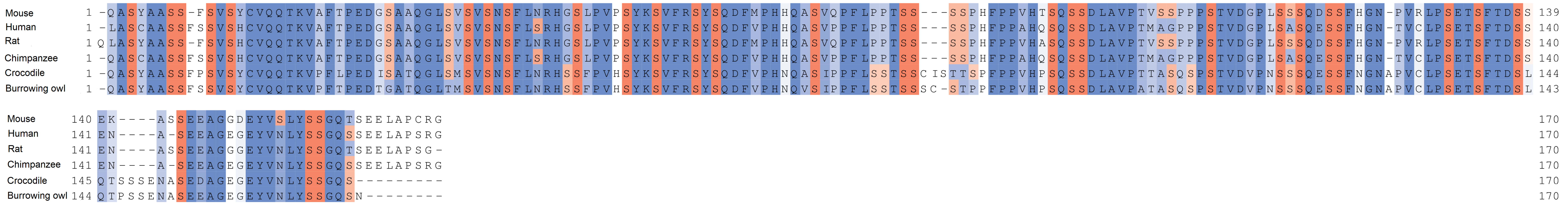


Fig. S5. Cassette exon encoded domain is serine-rich, and conserved across species. Mouse RAPGEF1 exons 12, 13, & 14 encode 170 amino acids inserted immediately following the fifth poly-proline tract. The totally and partially conserved amino acids are shown in dark and light shades of blue. Serine residues are marked in orange.

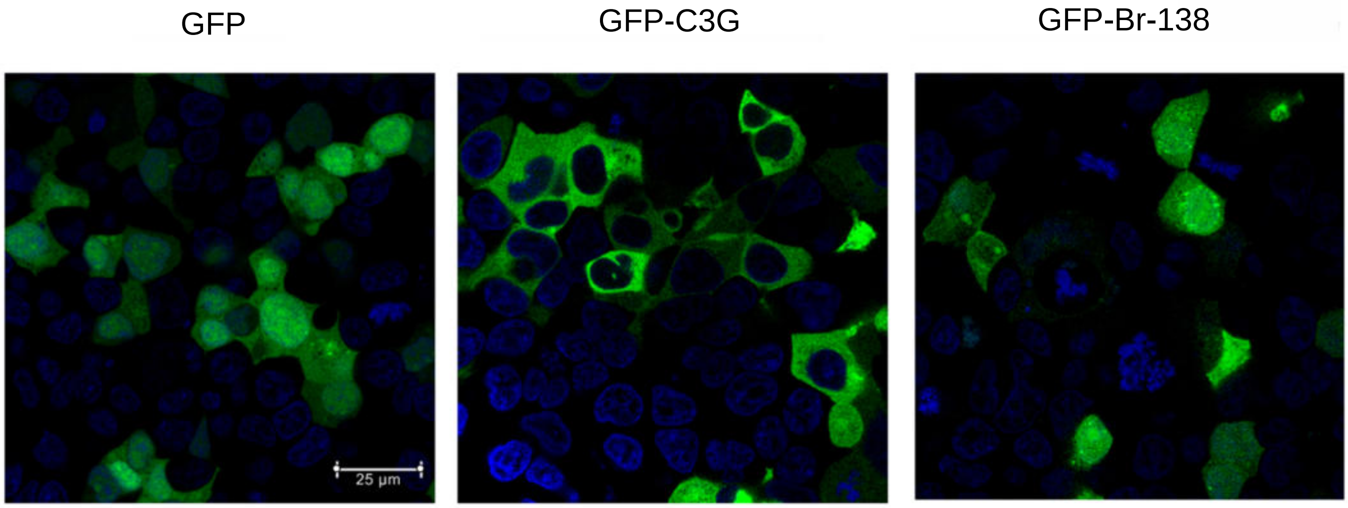


Fig. S6. The 138 aa insert in isoform 1 alters sub-cellular localisation of GFP. The amino acids encoded by exons 12 & 13 were cloned as a GFP-fusion protein using RNA isolated from the adult mouse brain (GFP-Br-138). Upon transient transfection in HEK293T cells, it concentrates in juxta-nuclear aggregates. GFP used as a control shows diffused staining throughout the cell, and RAPGEF1-GFP (full length isoform 3) shows predominant cytoplasmic staining.

Fig. S6

Table S1: Tissue specific isoform expression analyzed from RNA-seq data. Table showing expression of isoforms with various combinations of the cassette exons 12,13 & 14 in mouse tissues. Data was consolidated based on publicly available databases as indicated.

| **Transcript** | **ESCs** GSE159595 | **Brain** ENCLB532QIL | | **Skeletal muscle** GSM970865 | | **Testis** ENCLB555AYU | **Ovary** ENCSR000BZC | **Spleen** ENCLB809NWC | **Liver** ENCLB451FBJ | **Thymus** ENCLB888YNI |
|---|---|---|---|---|---|---|---|---|---|---|
| | | **E14.5** | **10 wks** | **P0** | **8 wks** | **8 wks** | **8 wks** | **8 wks** | **8 wks** | **8 wks** |
| NM_001039086.1 ENSMUST00000091146 | 2.67 | 0.96 | 4.12 | 1.6 | 9.79 | 0.13 | 0.48 | 5.98 | 0.69 | 19.35 |
| NM_001039087.1 ENSMUST00000095087 | 0 | 22.805 | 7.41 | 1.2 | 3.65 | 8.88 | 0.44 | 3.67 | 0 | 0 |
| NM_054050.2 ENSMUST00000102872 | 5.67 | 11.88 | 6.35 | 7.1 | 0.61 | 15.59 | 44.75 | 11.08 | 3.25 | 0 |
| NM_001362702.1 ENSMUST00000147755 | 0 | 18.26 | 7.51 | 0 | 2.75 | 10.46 | 0 | 2.91 | 1.38 | 0 |
| XM_006497592.4 ENSMUST00000238899 | 4.2 | 0 | 0 | 0 | 3.16 | 0 | 0 | 0 | 0 | 0 |
